# Fabrication and Characterization of Ceramic-Polymer composite 3D scaffolds and Demonstration of Osteoinductive propensity with gingival Mesenchymal Stem Cells

**DOI:** 10.1101/2023.03.20.533492

**Authors:** Manjushree M Bahir, Archana Rajendran, Deepak Pattanayak, Nibedita Lenka

**Author notes:** Authors for Correspondence: Nibedita Lenka, National Centre for Cell Science.

## Abstract

Bone tissue engineering involves the usage of metals, polymers, and ceramics as the base constituents in the fabrication of various biomaterial 3D scaffolds. Of late, the composite materials facilitating enhanced osteogenic differentiation/regeneration have been endorsed as the ideally suited bone grafts for addressing critical-sized bone defects. Here, we report the successful fabrication of 3D composite scaffolds with collagen type I (Col-I) in conjunction with three different crystalline phases of calcium-phosphate (CP) nanomaterials [hydroxyapatite (HAp), beta-tricalcium phosphate (βTCP), biphasic hydroxyapatite (βTCP-HAp or BCP)], obtained by altering the pH as the major variable. The fabricated 3D scaffolds consisting of ∼70 wt % CP nanomaterials and ∼ 30 Wt % of Col-I did mimic the ECM of bone tissue. The different Ca/P ratio and the orientation of CP nanomaterials in CP/Col-I composite scaffolds altered the microstructure, surface area, porosity, and mechanical strength of the scaffolds and also influenced the bioactivity, biocompatibility, and osteogenic differentiation of gingival-derived mesenchymal stem cells (gMSCs). The microstructure of CP/Col-I 3D scaffolds assessed by Micro-CT analysis revealed randomly oriented interconnected pores with pore sizes ranging from 80-250, 125-380, and 100-450µm respectively for βTCP/Col-I, BCP/Col-I, and HAp/Col-I scaffolds. Among these, the BCP/Col-I achieved the highest surface area (∼ 42.6 m^2^/g) and porosity (∼85%), demonstrated improved bioactivity and biocompatibility, and promoted osteogenic differentiation of gMSCs. Interestingly, the Ca^2+^ ions (3 mM) released from scaffolds could also facilitate the osteocyte differentiation of gMSCs *sans* osteoinduction. Collectively, our study has demonstrated the ECM mimicking biphasic CP/Col-I 3D scaffold as an ideally suited tissue-engineered bone graft.

## 1. Introduction

Bone defects and bone loss that affect the health quality of millions of people worldwide are majorly due to accidents and diseases like osteoporosis, cancer, infection, etc. The available treatment modalities for accident victims include the usage of allografts and autografts at large fracture edges. Unfortunately, several limitations of these grafts as well as the high demand-to-supply ratio of transplantable organs pose a major hindrance in this regard. Hence, attempts have been made to develop alternative bio-engineered bone grafts by fabricating artificial 3D scaffolds with the combination of cells to treat large-sized bone defects.^1-4^ Bio-nanomaterial mimicking the function of the native extracellular bone matrix has significant potential in bone tissue engineering. Hydroxyapatite (HAp) is a well-known calcium-phosphate (CP) nanomaterial and is the major inorganic component of the ECM composition of the bone matrix. Hence, CP-based nanomaterials hold unique chemical and structural properties similar to that of inorganic bone composition. Moreover, CP nanomaterials also possess excellent bioactivity, biocompatibility, osteoconductive, osteoinductive, and bio-degradability characteristics.^5,6^ Nevertheless, calcium-deficient HAp, such as di-, tri-, and octa-CP can also endure bone resorption, thereby having a substantial influence on the bone formation process.^7,8^ For instance, *in vivo* bone formation, involves the deposition of organic matrix and its successive mineralization. However, homeostasis between bone resorption and bone formation is essential for the osseous tissue establishment and bone remodelling activity.^9,10^

The fabrication of an ideal synthetic bone graft would depend on the composition of graft material that should be biocompatible and provide 3D stature, supporting cell adhesion and bone tissue growth as well as regeneration. This will have biological relevance mimicking *in vivo* conditions compared to that of a 2D substratum.^11,12^ In fact, several CPs such as HAp, beta-tricalcium phosphate (βTCP), biphasic CP, di-CP in combination with polymers like chitosan, polylactic acid (PLA), polycaprolactone (PCL), poly-L-lactide (PLLA), silk, alginate, etc. have been explored to design bioactive, bioresorbable, osteoconductive and osteoinductive 3D scaffolds to promote bone regeneration both *in vitro* and *in vivo*.^13-18^ Since the ECM composite of the native bone itself consists of collagen and CP material, 3D scaffolds containing these are of great interest in bioengineered bone tissue constructs. For example, collagen coating on titanium implants was reported to improve cell attachment and osteointegration potential of graft.^19,20^ Similarly, apatite/collagen composite with polymers such as PLLA and poly(D.L,lactide-co-glycolide) (PLGA) scaffolds also exhibited osteoblast-cell adhesion.^21,22^ However, *in vitro* seeding of mesenchymal stem cells (MSCs) on synthetic scaffolds to mimic the physicochemical microenvironment of bone is a prospective direction in bone tissue engineering.^23^

MSCs that are derived from various sources such as bone marrow, umbilical cord, placenta, periodontal ligament, tendon, etc. carry immunomodulatory properties and are bestowed with the intrinsic potential of giving rise to osteo-, adipo-, and chondrocytes.^24,25^ Hence, they can serve as an ideal cellular source for fabricating both auto- and allografts for bone tissue repair and regeneration, thereby having immense clinical implications. In fact, a number of reports do indicate the usage of MSCs to test the potential of bone graft scaffolds made up of various biomaterials, such as CP-ceramics, metals, and polymers.^26,27^ Efforts have also been made to understand the response and osteogenic differentiation capability of MSCs with different compositions of biomaterials. Indeed, the utilization of a CP-based porous 3D scaffold can offer an appropriate milieu for cell-matrix interaction. Moreover, the viability, adhesion, proliferation, and differentiation of MSCs to various lineages are affected not only by the component of composite materials but also by the microstructure, porosity, and surface area of the scaffolds. Additionally, bioactive grafts are required to interact with the surrounding bone-forming tissue.^28,29^ In this regard, the ideal strategy would be to incorporate the organic/inorganic composite biomaterial scaffolds, which would mimic ECM of bone composition and thereby promote osteointegration potential. The dexamethasone-loaded CP/collagen composite 3D scaffold has been shown to promote osteogenic differentiation of human MSC both *in vitro* and *in vivo*.^30,31^ Chen et al. have also demonstrated that the HAp-collagen coating on decellularized bone favoured MSCs adhesion, proliferation, and their osteogenic differentiation potential.^32^

In this study, we have reported the fabrication of ceramic-polymer composite scaffolds, namely HAp/Col-I, βTCP/Col-I, and βTCP-HAp (BCP)/Col-I by freeze-drying technology followed by UV cross-linking. The stated three different phases of CP nanomaterials, i.e., HAp, βTCP, and BCP have been achieved by the wet chemical method by keeping the pH as the sole variable. Assessment of physicochemical characteristics of thus developed scaffolds revealed those to retain the functional groups of CP nanomaterials and possess an interconnecting porous structure with decent porosity and mechanical strength. Moreover, the fabricated composite scaffolds were not only found to be bioactive but also promoted osteogenic differentiation of gingival tissue-derived MSCs (gMSCs) under maintenance conditions itself. Further investigations also revealed their suitability for bone tissue engineering applications by seeding gMSCs onto those and subjecting them to osteoinduction. Interestingly, BCP/Col-I was found to be the most potent composite scaffold among the three, thereby offering the potential utility of the same in stem cell-based therapy for bone regeneration.

## 2. Materials and methods

### 2.1. Synthesis of CP nanomaterial

Different phases of CP nanomaterials were synthesized by the wet-chemical precipitation method by using calcium nitrate tetra-hydrate (CaNO_3_.4H_2_O) and di-ammonium hydrogen orthophosphate (NH_4_)_2_HPO_4_) as the source of Ca and P respectively.^33^ Briefly, 0.6M di-ammonium hydrogen orthophosphate was added drop-wise to the 1M calcium nitrate tetra-hydrate solution at a rate of 1ml/min resulting in a milky precipitate. The pH of the precipitate was adjusted to 8, 9, and 11 using liquid ammonia to get different phases of CP nanomaterials. The reaction was carried out at 70 °C under vigorous stirring for 4 h followed by 24 h aging at room temperature and the precipitate was washed several times with double distilled water to remove the residual ammonia. Further, the resultant precipitate was poured into the petridish for drying at room temperature for 2 days followed by sintering at 1000 °C for 2 h using a muffle furnace with a heating rate of 5 °C/min. The calcined CP nanopowders were characterized by Fourier Transform Infrared (FT-IR) spectra for functional group analysis and X-ray diffractometer (XRD) for pH-dependent phase formation (HAp, βTCP, and βCP) and crystal structure determination.

### 2.2. Fabrication of 3D Scaffold

Col-I from bovine achilles tendon (Sigma Aldrich) was dispersed in 0.05M glacial acetic acid, and mixed with each of the afore-stated CP nanopowders independently followed by their freezing at -80 °C for 24 h. The CP/Col-I (70:30 wt %) scaffolds were fabricated by freeze-drying using a lyophilizer (C-gen Biotech Ltd) maintained at -80 °C and with 1 Pa pressure to obtain HAp/Col-I, βTCP/Col-I, and BCP/Col-I ceramic-polymer composite scaffolds. Further, UV crosslinking was carried out to crosslink Col-I, as described.^34^ Subsequently, the scaffolds were washed thoroughly using deionized water to remove any trace amount of glacial acetic acid followed by freeze-drying.

### 2.3. Characterizations of 3D scaffolds

#### 2.3.1. X-ray diffraction (XRD) and FT-IR analysis

The fabricated 3D scaffolds were characterized by XRD to confirm the retention of different phases of CP as that noted with parental nanomaterials (HAp, βTCP, and BCP). The XRD spectra were recorded on an X-ray diffractometer (XRD, XPERT-PRO) using CuKα (1.54 Å) radiation over 2θ range between 20 to 60° with a step size of 0.02°. Further, the functional group analysis of fabricated 3D scaffolds (HAp/Col-I, βTCP/Col-I, and BCP/Col-I) was carried out by FT-IR spectra between 500-4000 cm^-1^ region on Bruker Optics ALPHA-E spectrometer using a Diamond ATR (attenuated total reflection) (Golden Gate) with a resolution of 1cm^-1^. The crystallite size was calculated from the XRD by the Debye-Scherrer equation [D = (kλ/β cos θ)].

#### 2.3.2. Transmission Electron Microscope

The size, shape, and crystal nature of CaP powder synthesized at pH 8, 9 and 11, calcined at 1000 ⁰C were observed under a transmission electron microscope (TEM; Tecnai 20 G2 FEI, The Netherlands). The specimen for TEM observation was prepared from the particle dispersed in ultrapure water. A drop of the dispersed particles was placed on the copper grid and allowed to dry before observation.

#### 2.3.3. Field emission scanning electron microscopy (FE-SEM) and energy dispersive X-ray (EDAX) analysis

The microstructure and elemental analysis of developed 3D scaffolds were carried out using FE-SEM (Carl-Zeiss, Germany) coupled with an energy dispersive spectrometer. Before imaging, the 3D scaffold surface was sputter-coated with gold to avoid the surface charging effect, and the images were acquired at various magnifications after placing the scaffolds on the FE-SEM sample holder. For elemental analysis through EDAX, the scaffold surface was scanned, and the quantitative information was obtained by considering the spectral peaks.

#### 2.3.4. Micro-Computed Tomography (micro-CT) Analysis of CP/Col-I 3D scaffolds

The internal micro-structure and the existence of the interconnected pores within the 3D scaffolds were analysed by micro-CT. The CP/Col-I 3D scaffolds were imaged by X-Ray Micro-CT scanner (SkyScan 1276; Bruker micro-CT) (acquisition parameters settings: source voltage = 50 kV; source current = 200 μA; rotation step = 0.6°; rotation angle = 180°; 0.5 mm aluminium filter; Image Pixel Size = 7 μm; phantom = HAp). The software NRecon (Version: 1.0.13; Bruker micro-CT), was used to reconstruct the cross-section images of the 3D scaffolds, and the application 3D Creator (CTAn software, Bruker micro-CT,) was used for the reconstruction of the 3D model. The pore size of the 3D scaffolds was analysed using the binary image dataset and sections of 2D transverse planes.

#### 2.3.5. Assessment of mechanical strength

The tensile properties of the HAp/Col-I, βTCP/Col-I, and BCP/Col-I 3D scaffolds were determined using a Universal Testing Machine (Model STS 248; Star testing system, India). Briefly, the 3D scaffolds were cut into rectangular strips (2.5 cm × 1 cm) with a thickness of 1 mm. Each end of the scaffold strip was placed in a metal clamp mounted on an STS 248 against the applied cell load of 980 N with a crosshead speed of 10 mm/min. The maximum tensile strength and elongation before fracture were determined at room temperature. The experiments were performed in triplicate and the average values were reported for the mechanical (tensile) strength test.

#### 2.3.6. Analysis of specific surface area and porosity

To determine the specific surface area of the 3D scaffold nanocomposites, their N_2_ adsorption-desorption isotherms were recorded at 77 K in a Quantachrome®ASiQwin (Autosorb iQ Station 1). The liquid substitution method was used to analyse the open porosity of 3D scaffolds by taking a known volume of ethanol (V1) in a graduated cylinder and immersing each scaffold with identical weight in it, as described.^35^ The increase in the volume of ethanol in each after 5 min of immersion was considered as V2. Further, after removing the scaffold, the residual ethanol volume was measured and considered as V3. The porosity of the 3D scaffold was calculated by using the formula:^35^ [Porosity % = [(V1−V3)/(V2−V3)] ×100].

### 2.4. Evaluation of 3D scaffolds for bone tissue engineering applications

#### 2.4.1. Determination of Bioactivity of 3D scaffolds

*In vitro* bioactivity test of HAp/Col-I, βTCP/Col-I, and BCP/Col-I 3D scaffolds were performed using simulated body fluid (SBF) prepared using the protocol described.^33^ The scaffolds were incubated with SBF using a CO2 incubator maintained at 37 °C and with 5% CO2. Two different time points (8- and 32 days) were considered for the bioactivity assessment with replenishing SBF every 3 days. At the end of predetermined immersion durations, 3D scaffolds were removed from SBF and rinsed with distilled water followed by drying at room temperature. Further, surface morphology and elemental analysis of 3D scaffolds were carried out by FE-SEM and EDAX respectively, as described before.

#### 2.4.2. Evaluation of biocompatibility of 3D scaffolds

The gMSCs already available in the laboratory were considered for the assessment of the biocompatibility of the stated scaffolds by MTT [3-(4,5-dimethylthiazol-2-yl)-2,5-diphenyltetrazolium bromide] colorimetric assay using the standard protocol.^27^ Briefly, 1 x 10^4^ cells were seeded on HAp/Col-I, βTCP/Col-I, and BCP/Col-I composite 3D scaffolds and were cultured in a humidified CO2 incubator at 37 °C using Dulbecco’s modified Eagle medium (DMEM) supplemented with 10% FBS, 2mM L-glutamine, and Penicillin-Streptomycin as the maintenance medium. After 72 h, the culture medium was removed gently from each well and the cell-laden scaffolds were incubated with MTT solution for 4 h to form formazan crystals in cells. Further, formed purple formazan crystals were solubilized by adding dimethyl sulfoxide (DMSO) and incubating it for 1 h. Finally, absorbance was measured at 570 nm using a microplate reader (Thermo multiskan GO) for quantifying the metabolic activity of viable cells. The gMSCs maintained without the scaffolds were considered as control (Ctrl) and the % viability was assessed by keeping the Ctrl value as 100%. Further, to monitor the infiltration, adhesion and growth of gMSCs post-seeding, the gMSCs-laden scaffolds were subjected to immunostaining to detect the cellular response in terms of effective cell adhesion and growth after 72 h of their maintenance, as described.^27^

#### 2.4.3. Evaluation of osteogenic differentiation in gMSCs-laden scaffolds in vitro

To evaluate the osteogenic potential of gMSCs, 1 x 10^4^ cells were seeded on CP/Col-I scaffolds and cultured for 72 h using DMEM, as described in the previous section. The culture medium was replaced with an osteoinduction medium (Invitrogen) to induce osteogenic differentiation with medium replenishment done every 3 days. In parallel, gMSCs maintained on tissue culture dishes without any scaffold and both with and without osteoinduction conditions were considered as positive and negative controls respectively. On day 21, all the groups were stained with Alizarin red-S (ARS) staining to confirm the deposition of the mineralized matrix and to confirm the osteogenic differentiation of gMSCs. The scaffolds were also maintained under the maintenance medium and processed in a similar way to monitor the background, if any. While the quality of osteocytes-induced mineralization was determined under an inverted microscope (Nikon) by monitoring ARS staining, spectrophotometry was carried out to quantify the deposited OCN by incubating the samples in 10% glacial acetic acid at room temperature for 30 min and measuring the absorbance at 405 nm using a plate reader.^27^ The OD in the case of cell-laden scaffolds was normalized with respect to each scaffold background and represented as the relative absorbance unit (AU).

#### 2.4.5. Determination of Calcium Ion release

Release of calcium ion from HAp/Col-I, βTCP/Col-I, and BCP/Col-I nanocomposite scaffolds was carried out by incubating the same in PBS with continuous stirring at 100 RPM and at 37 °C. Further, the release of Ca^2+^ ion was estimated at various time intervals (3-, 7- 15-, and 21 days) by ethylene-di-amine-tetra-acetic acid (EDTA) titration method.^36^

### 2.5. Statistical analysis

The experiments were carried out in triplicates and repeating for 3-4 times each. All quantitative data were presented as mean ± standard error of the mean (SEM). Statistical comparison was performed using t-test/paired t-test (Sigmaplot) analysis and the statistical difference was equated at the p-value. [P ≤0.05 represented as one star (*), P ≤0.01 two stars (**), P ≤0.001 three stars (***)].

## 3. Results

### 3.1. Fabrication and Characterization of CP/Col-I 3D scaffolds

#### 3.1.1. Structural analysis by XRD

Synthesis of different phases of CP nanomaterials by wet chemical method primarily depends upon various parameters, such as; the initial precursor Ca/P molar ratio, reaction temperature, pH, stirring rate, and sintering temperature.^37^ The phase analysis of CP nanomaterials utilized for the fabrication of CP/Col-I scaffold was confirmed from XRD spectra shown in Fig. 1a. The XRD spectra of synthesized CP nanomaterials were compared with the JCPDS data no. 09-0432 for HAp and JCPDS. 09-0169 for the βTCP phase. The maximum diffraction peak intensity at 211 plane in the synthesized CP nanomaterial at pH-11 indicated the development of HAp hexagonal phase. In fact, the absence of any other phase formation could attest to the purity of HAp. Moreover, XRD spectra of CP nanomaterial synthesized at pH-8 confirmed the βTCP phase formation. Similarly, the BCP synthesized at pH-9 could indicate the presence of ∼27% of βTCP and ∼73% HAp phases. The presence of the secondary βTCP phase in BCP was further confirmed by overlapping the XRD peak profiles of HAp and βTCP materials where the relative intensities of a diffraction peak resided at 2θ = 31.9 ° corresponding to HAp, and a minor intensity diffraction peak at 2θ = 31.2 ° represented βTCP phase.

**Fig. 1:**
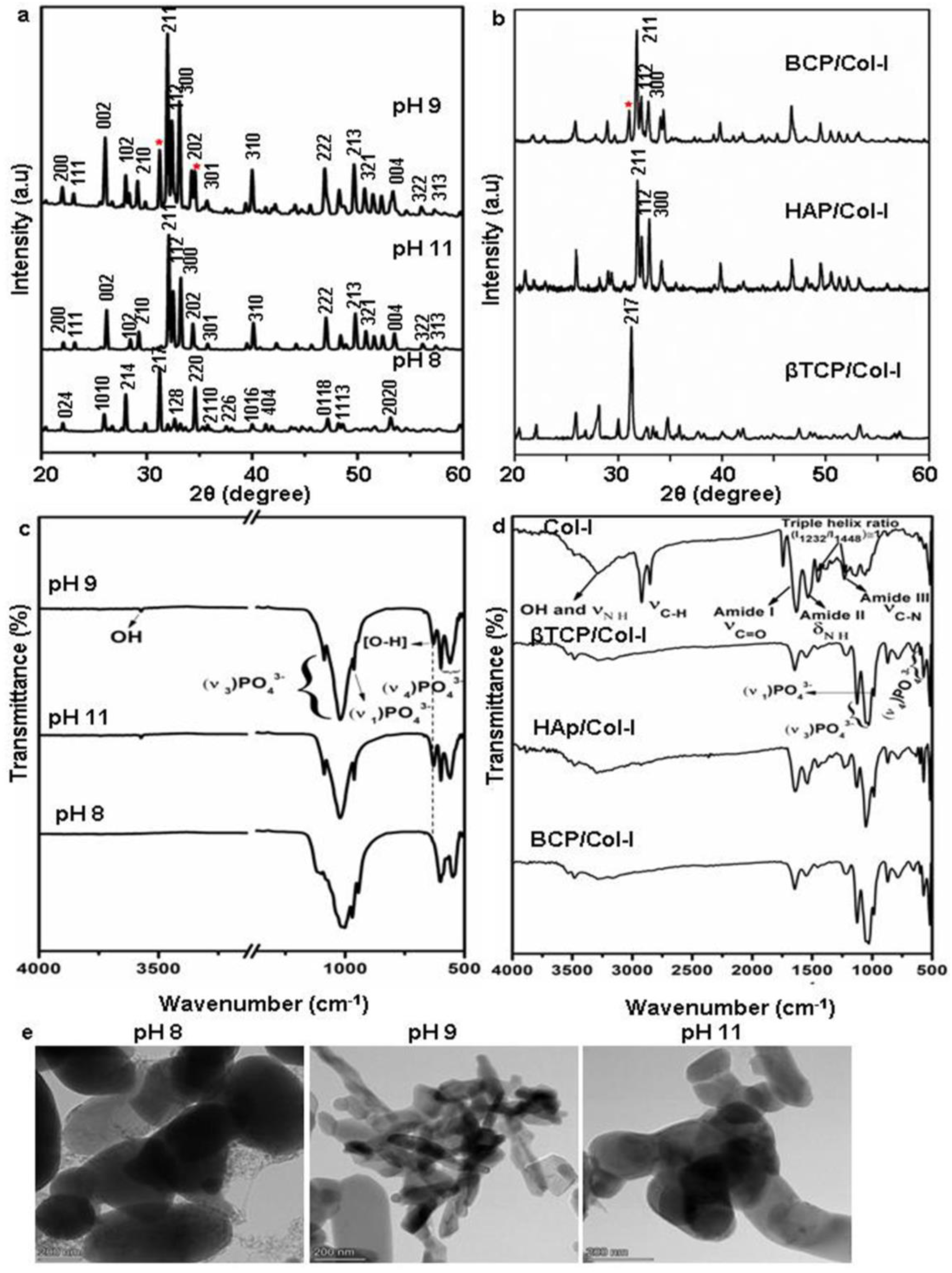
(a, b) X-ray diffraction pattern of (a) CP nanomaterials synthesized by wet-chemical precipitation method at PH 8, 9, and 11 and (b) CP/Col-I 3D scaffolds fabricated by freeze drying technique (* indicates the presence of secondary βTCP in HAp structure). (c, d) FTIR spectra of CP synthesized nanomaterials (c) and CP/Col-I 3D scaffolds (d). (e) TEM images of CP nanomaterials synthesized at pH 8, 9, and 11 by wet-chemical precipitation method. Scale in ‘c’ has been adjusted to show the OH- (3492 cm−1) peak.

The architecture of a 3D scaffold is of more biological relevance since it can support the cells in 3D and mimic the *in vivo* scenario.^12^ Fig. 1b illustrates the XRD pattern of the CP/Col-I scaffold, fabricated by the freeze-drying technique. The main XRD peak position associated with the βTCP phase was noted at 2θ = 31.2 ° in βTCP/Col-I scaffold, 2θ = 31.9 ° for HAp in HAp/Col-I and 2θ = 31.2 and 31.9 ° for βTCP-HAp in BCP/Col-I and validated the association of crystalline CP nanomaterials with the Col-I composite. The crystallite size was 39 nm, 35 nm, and 70 nm respectively for HAp, βTCP, and BCP nanoparticles as discerned by using by Debye-Scherrer equation. However, the broadened XRD peaks of CP nanomaterials in the CP/Col-I scaffolds suggested the reduction in the crystallinity of CP due to its conjugation with Col-1 in the matrix composite.

#### 3.1.2. Functional group analysis by FTIR

The crystal structure of HAp [Ca_10_ (PO_4_)_6_(OH)_2_] involves phosphate (PO_4_^3-^), hydroxyl (OH^-^) functional groups, whereas βTCP [Ca_3_(PO_4_)_2_] involves only (PO_4_^3-^) functional group. The FTIR spectra of CP nanomaterials synthesized at pH 8, 9, and 11 respectively are seen in Fig. S1 (Original) and Fig. 1c (Scale adjusted). The characteristic absorbtion bands for PO_4_^3-^ and OH^-^ were observed in the 4000 -500 cm^-1^ IR range. The PO_4_^3-^ absorbtion bands at 562 cm^-1^, 602 cm^-1^ corresponding to asymmetric bending vibration (υ_4_), 961 cm^−1^, 1032 cm^-1^ to symmetric (υ1), and 1091 cm^-1^ to asymmetric stretching vibration (υ3) modes were identified in all the CP nanomaterials synthesized at different pH. However, an additional very low intense shoulder absorbtion peak for PO_4_^3-^ at 945 cm^-1^ was seen in the case of BCP. Although, the existence of bands at 3573 cm^−1^ and 631 cm^−1^ due to the stretching vibration and liberation mode of the OH^-^ group were clearly seen in CP nanomaterials synthesized at both pH-9 and -11, the peak intensity in BCP was comparatively low, hence supporting the biphasic phase formation. Furthermore, the disappearance of the stated absorption bands for OH^-^ groups in the IR spectra represented the lack of OH^-^ group in the CP nanomaterial synthesized at pH-8. Taken together, the structural and chemical analysis of CP nanomaterials synthesized by wet chemical method with varying pH at 8, 9, and 11 aligned well with the βTCP, BCP, and HAp phases respectively.^38-40^ Hence, this attested to the authenticity of our fabrication approach.

Fig. 1d shows the FTIR spectra of Col-I and CP/Col-I 3D composite scaffolds. The IR spectra of Col-I depicting the absorbtion bands at 1634 cm^-1^, 1540 cm^-1^, and 1240 cm^-1^ were corresponding to the amide I, amide II, and amide III respectively. Moreover, the absorbtion bands at 1450 cm^-1^, 1397 cm^-1^, and 1338 cm^-1^ corresponding to δas CH_3_ side-chain vibrations and 3310 cm^-1^ (-OH), 3310 cm^-1^ (N-H) belonging to amide A, amide B were observed. The presence of absorbtion bands corresponding to PO_4_^3-^ (1100 cm^−1^, 1095 cm^-1^, 1035 cm^-1^, 970 cm^-1^, 950 cm^−1^, 605 cm^−1^, and 570 cm^-1^), OH^-^ (3492 cm^−1^ and 630 cm^−1^) and amide functional groups from Col-I in CP/Col-I composite scaffolds confirmed the cross-linking between the Col-I and CP nanomaterials. Further, the presence of a carbonate peak at 870 cm^-1^ observed in IR spectra in all 3D scaffolds revealed the appearance of a carbonate group during scaffold composite preparation.

#### 3.1.3. Microstructural and elemental analysis

The size and shape of the CaP particles synthesized at various pH (8, 9, and 11) and calcined at 1000 °C were studied using TEM. As seen in Fig. 1e, the particle size of CaP was in the range of 500 nm -1μm, irrespective of the pH used. While relatively globular-shaped particles were observed in the case of CaP synthesized at pH 8, the micrographs revealed a majority having rod/elongated particles in the case of pH 11 synthesized ones. Interestingly, the CaP synthesized at pH 9 showed a mixture of both globular and elongated CaP particles. In line with XRD and FTIR spectra, the TEM micrographs also confirmed the pH-dependant mono- and biphasic patterns of HAp. The particles looked agglomerated in all the cases possibly due to high specific energy during calcination.

The porous microstructure of the scaffolds contributes to providing structural support for new tissue formation and space for cell adhesion, proliferation, and differentiation into desired tissue types. The surface morphology, porous architecture, and elemental analysis of the CP/Col-I scaffolds were investigated by FE-SEM/EDAX. The porous microstructure of βTCP/Col-I showed a homogeneous distribution of leaf-like structure with the typical size of 11 ±0.5 μm, whereas petal-like morphology was noticed on the backbone of HAp/Col-I and BCP/Col-I 3D scaffolds (Fig. 2a-f). Together, these findings demonstrated the different orientations of CP nanomaterials with Col-I. The higher resolution FE-SEM images of the CP/Col-I scaffolds shown in Fig. 2d-f revealed uniform dispersion of Col-I with HAp and BCP nanomaterials confirming their higher affinity with the Col-I unlike that of βTCP. Interestingly, the BCP/Col-I scaffold presented a porous microstructure with increased roughness on the surface and hence, portrayed having increased surface area for better cell adhesion, proliferation, migration, tissue-specific growth, and nutrient flow compared to βTCP/Col-I and HAp/Col-I scaffolds. Further, the EDAX analyses revealed that the peaks corresponding to Ca, P, and O elements were present in CP/Col-I scaffolds (Fig. 2g-i). The Ca/P atomic ratios in βTCP/Col-I, BCP/Col-I, and HAp/Col-I scaffolds were observed to be 1.1±0.1, 1.5±0.05 and 1.67±0.02 respectively, which were in good agreement with XRD results.

**Fig. 2:**
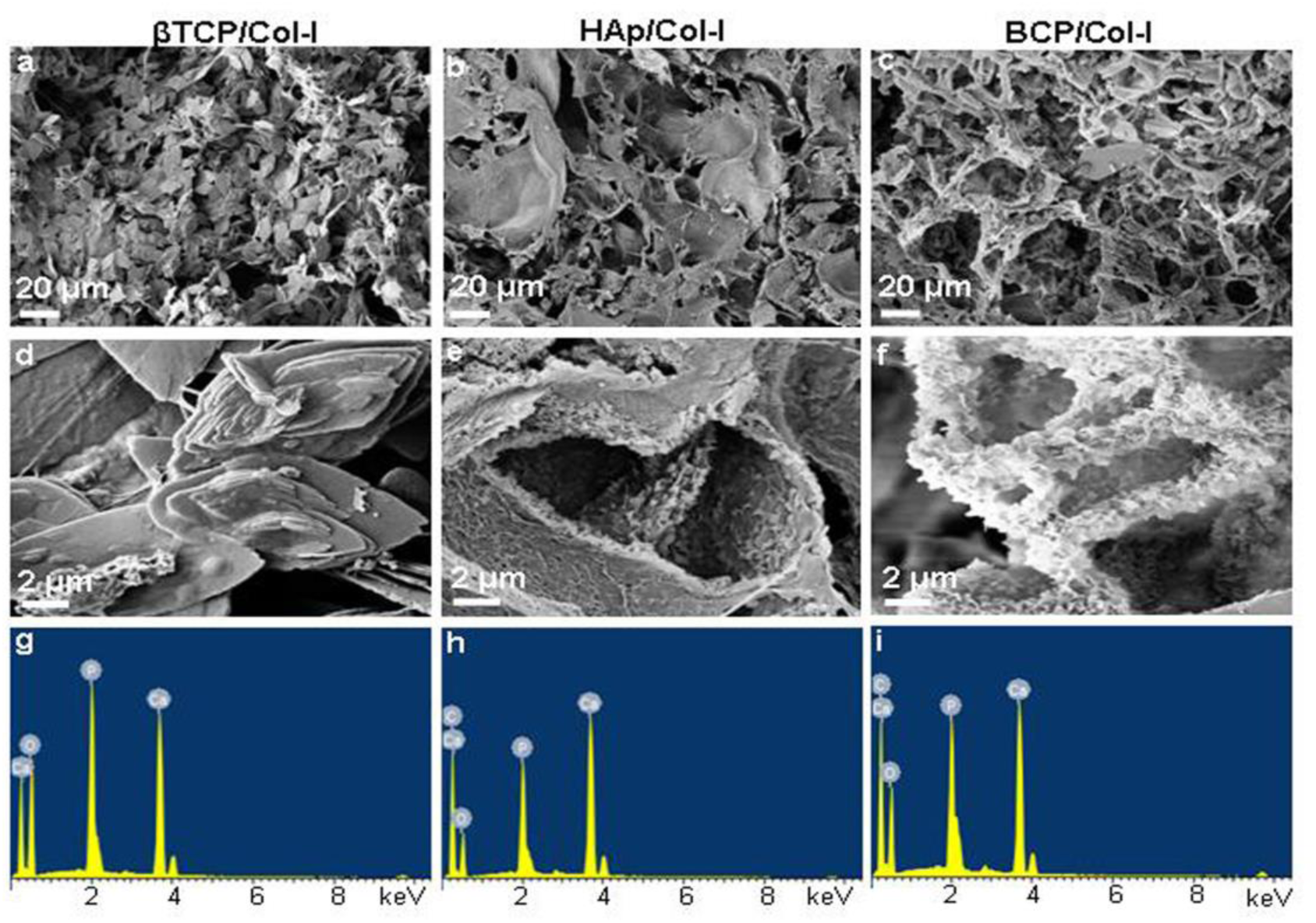
FE-SEM images (a-f) and EDAX spectra (g-i) of CP/Col-I 3D scaffolds developed by freeze drying technique. The microstructure of βTCP/Col-I (a,d) showed a homogeneous distribution of leaf-like structure with the typical size of 11 ±0.5 μm; petal-like morphology was observed on the backbone of HAp/Col-I (b,e), and BCP/Col-I (c,f) 3D scaffolds; (a-c) 1.5 KX magnification and (d-f) 25 KX magnification. EDAX spectra (g-i) of βTCP/Col-I, HAp/Col-I, and BCP/Col-I scaffolds showed the presence of Ca, P, and O elements in CP/Col-I scaffolds.

#### 3.1.4. Porosity and surface area

The porous biomaterial scaffolds with higher surface area offers adsorption of bone morphogenic protein which can help to promote the bone regeneration process. Additionally, the 3D CP-scaffold with porous structure with interconnected micro-pores enables better cell adhesion, migration, and growth thereby facilitating integration into host bone and enhanced osteoinduction following grafting.^32^ Fig. 3a displays the 2D (i-vi) and 3D (vii-ix) micro-CT images of CP/Col-I 3D scaffolds. Micro-CT images of CP/Col-I 3D scaffolds revealed randomly sized well-interconnected pores with pore sizes ranging from 80-250µm for βTCP/Col-I, 125-380 µm for BCP/Col-I and 100-450 µm for HAp/Col-I scaffold. The pore size was obtained from the analysis of the 2D transverse plane of the scaffolds, as shown in Fig. 3a (i-iii). Furthermore, the porous architecture with interconnected pores of CP/Col-I 3D scaffolds was also confirmed from the 2D images of the coronal plane (iv-vi) and 3D (vii-ix) images of the scaffolds. Similarly, the average porosity in the fabricated scaffolds analysed by the liquid displacement method was observed to be 65±0.75%, 75±0.28%, and 85.76±1.33% for βTCP/Col-I, HAp/Col-I, and BCP/Col-I 3D scaffolds respectively (Fig. 3b). In fact, that complied well with the porosity usually seen in case of cancellous bones.^26^ Our findings indicated that the porous BCP/Col-I composite possessed a higher surface area (42.6m2/g) compared to HAp/Col-I (35.6m2/g) and βTCP/Col-I (35.7m2/g) scaffolds (Fig. 3c). The surface area, pore-volume and average pore size of the CP/Col-I scaffolds analysed by BET has been shown in Table-1. Overall, the CP/Col-I scaffolds exhibited type-IV adsorption/desorption isotherms representing both micro- and mesoporous scaffold architecture.

**Fig. 3:**
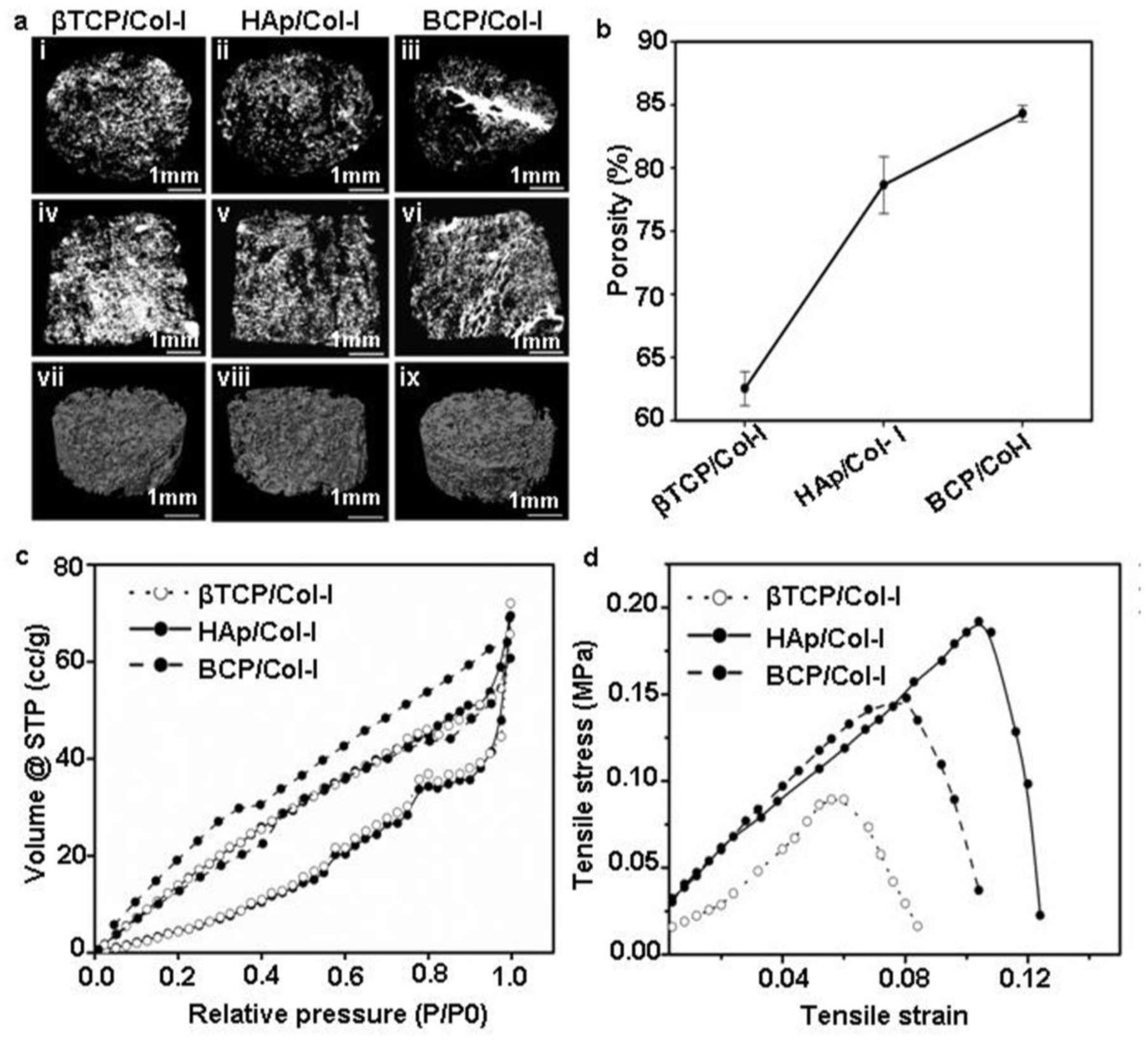
(a) Micro-CT analysis of the CP/Col-I 3D scaffolds. (i-iii) the transverse plane, (iv-vi) coronal plane, and (vii-ix) 3D reconstruction images of the scaffolds demonstrating the porous architecture with interconnected pores. The pore size ranged between 80-250 μm for βTCP/Col-I, 125-380 μm for BCP/Col-I, and 100-450 μm for HAp/Col-I scaffolds [scale bar: 1 mm (i-vi) and 200 μm (vii-ix)]. (b) Porosity of CP/Col-I scaffolds was determined by the liquid displacement method showing maximum porosity with BCP/Col-I followed by HAp/Col-I and βTCP/Col-I. (c) Surface area of CP/Col-I 3D scaffolds was determined by the BET method based on adsorption and desorption of N2 revealing comparatively higher surface area (42.6m2/g) with BCP/Col-I scaffold than that of HAp/Col-I (35.6m2/g) and βTCP/Col-I (35.7m2/g) scaffolds. (d) The ultimate tensile strength of CP/Col-I 3D scaffolds was observed in the range of 1.33 to 1.9 Mpa.

**Table-1:**
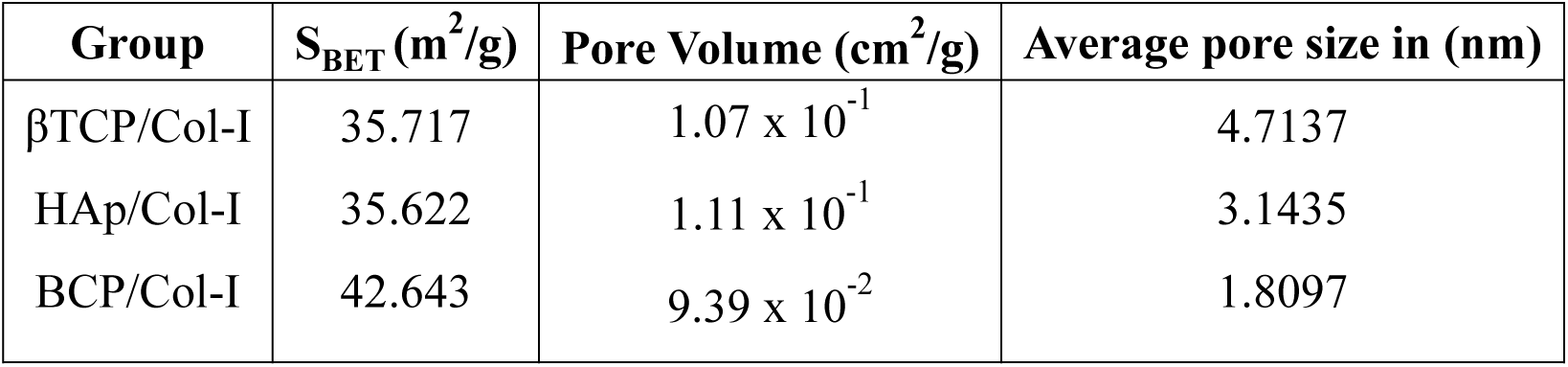
BET surface area, pore volume and pore size of CP/Col-I scaffolds.

#### 3.1.5. Mechanical strength analysis

The implantable scaffolds undergo substantial stress and strain in surgical intervention during bone grafting. As a consequence, requirement of ductile and flexible scaffold becomes effective carrying clinical significance.^41,42^ Fig. 3d shows the tensile stress-strain curves of the CP/Col I 3D scaffolds. The ultimate tensile strength was maximum in HAp/Col-I composite scaffold (1.9 MPa) followed by BCP/Col-I composite (1.7 MPa) and was the least in βTCP/Col-I (1.33 MPa) composite scaffold. Indeed, the tensile strength of CP/Col-I scaffolds fabricated by us matched well with the tensile strength of a trabecular bone (1-5 MPa).^43^ The reduction in ultimate tensile strength in βTCP/Col-I could be possibly due to the higher degradation/resorbtion rate of βTCP crystals and the difference in their interaction or orientation with Col-I fibrils, also supporting the microstructural analysis, as shown in Fig. 2.

### 3.2. *In vitro* biological properties of 3D scaffolds

#### 3.2.1. In vitro bioactivity

Bioactive performance of a 3D scaffold is dependent on its ability to allow the formation of newly calcified tissue on its surface called osteointegration. *In vitro* SBF test of synthetic bone graft is a standard method to envisage *in vivo* bone bonding or osteointegration ability.^44^ A newly formed apatite layer was nucleated on the surface of 3D scaffolds by forming cauliflower-like morphology after 8 days of SBF incubation, depicted in Fig. 4. Interestingly, the morphology of the newly formed apatite layer on the surface of all 3D scaffolds seemed rather similar after 8 days of incubation in SBF. However, the nucleation rate seemed to be superior on BCP/Col-I composite scaffold, whereas the least nucleation was seen in the case of βTCP/Col-I (Fig. 4a). On day 32 however, the nucleation of apatite on the surface of all the three CP/Col-I 3D scaffolds was greatly improved covering the entire surface to indicate a homogeneous nucleation occurring over a period (Fig. 4b). Further, the elemental analysis suggested that the newly formed apatite layer involved oxygen (O), phosphate (P) and calcium (Ca) ions (Fig. S2a, b). Together, these findings suggested that the introduction of ∼27% βTCP in HAp accelerated the bioactivity of the BCP/Col-I 3D scaffold.

**Fig. 4:**
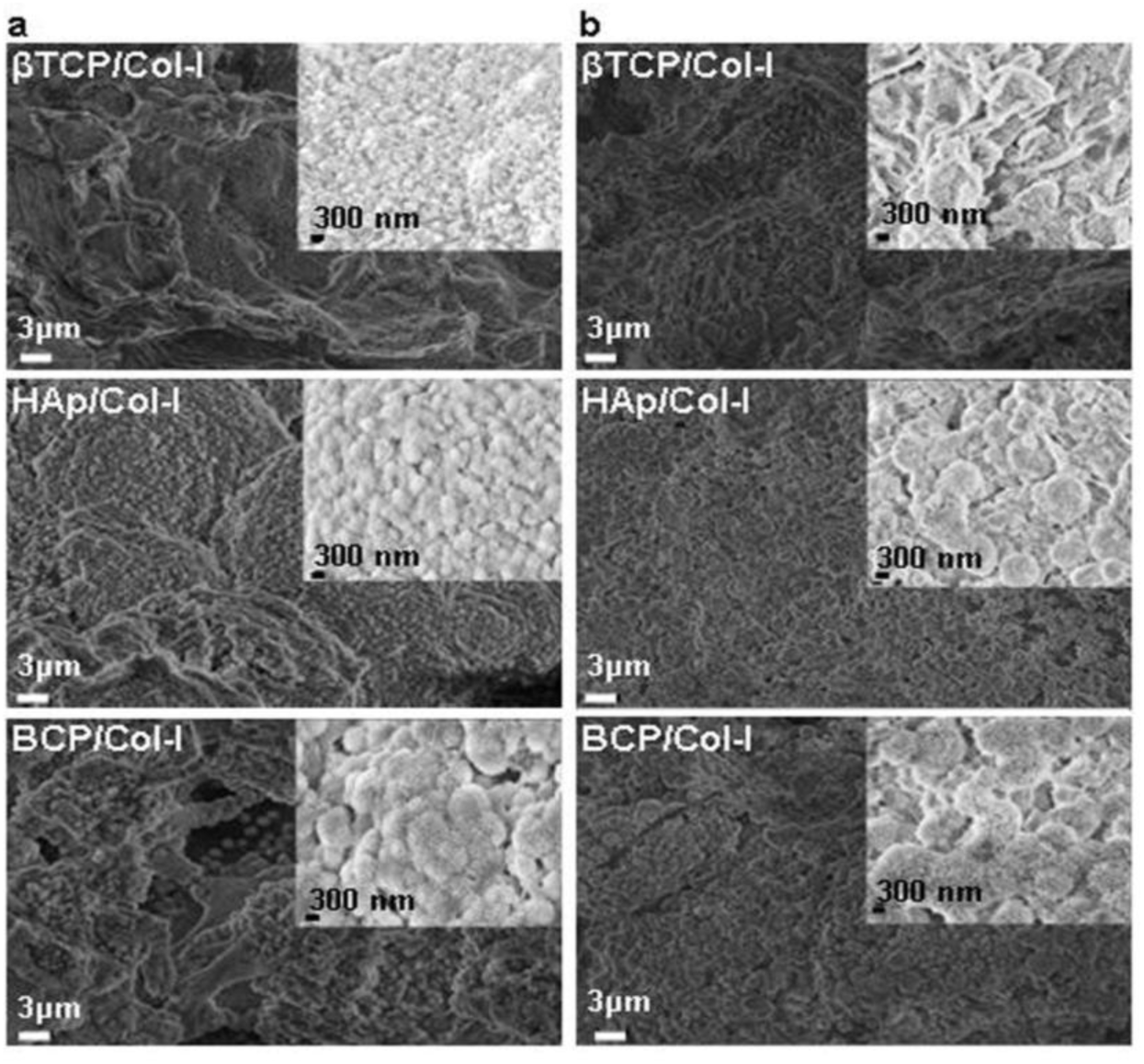
Bioactivity of the 3D scaffolds. FE-SEM images displaying apatite layer formation on the surface of CP/Col-I 3D scaffold after incubating in SBF for 8 days (a), and 32 days (b) time points respectively; inset shows the magnified (25 KX) images of apatite layer.

#### 3.2.2. *In vitro* biocompatibility of CP-Col-I 3D scaffolds

Considering the high demand-to-supply ratio of transplantable organs, the need of the time is to develop bioengineered organs. The seeding of human MSCs onto synthetic bone scaffolds is of great interest for the reconstruction of bone defects.^28^ Therefore, in this study, human gMSCs have been used to investigate the biocompatibility of the CP/Col-I scaffolds. *In vitro* biocompatibility of 3D scaffolds was evaluated by seeding gMSCs on scaffolds for a period of up to 3 days. Fig. 5a shows the gMSCs growing in presence of CP-Col-I scaffolds similar to that in control. Cytotoxicity of 3D scaffolds was assessed by means of MTT colorimetric assay. MTT results clearly demonstrated that both HAp-Col-I and BCP/Col-I scaffolds did not have a significant effect on the cell viability unlike that seen with βTCP/Col-I 3D scaffold (Fig. 5b). In fact, among these three composite scaffolds, BCP/Col-I showed superior biocompatibility. Nevertheless, the cell viability was observed to be > 80% in the case of all three CP/Col-I composite scaffolds, thereby attesting to their biocompatibility with gMSCs.

**Fig. 5:**
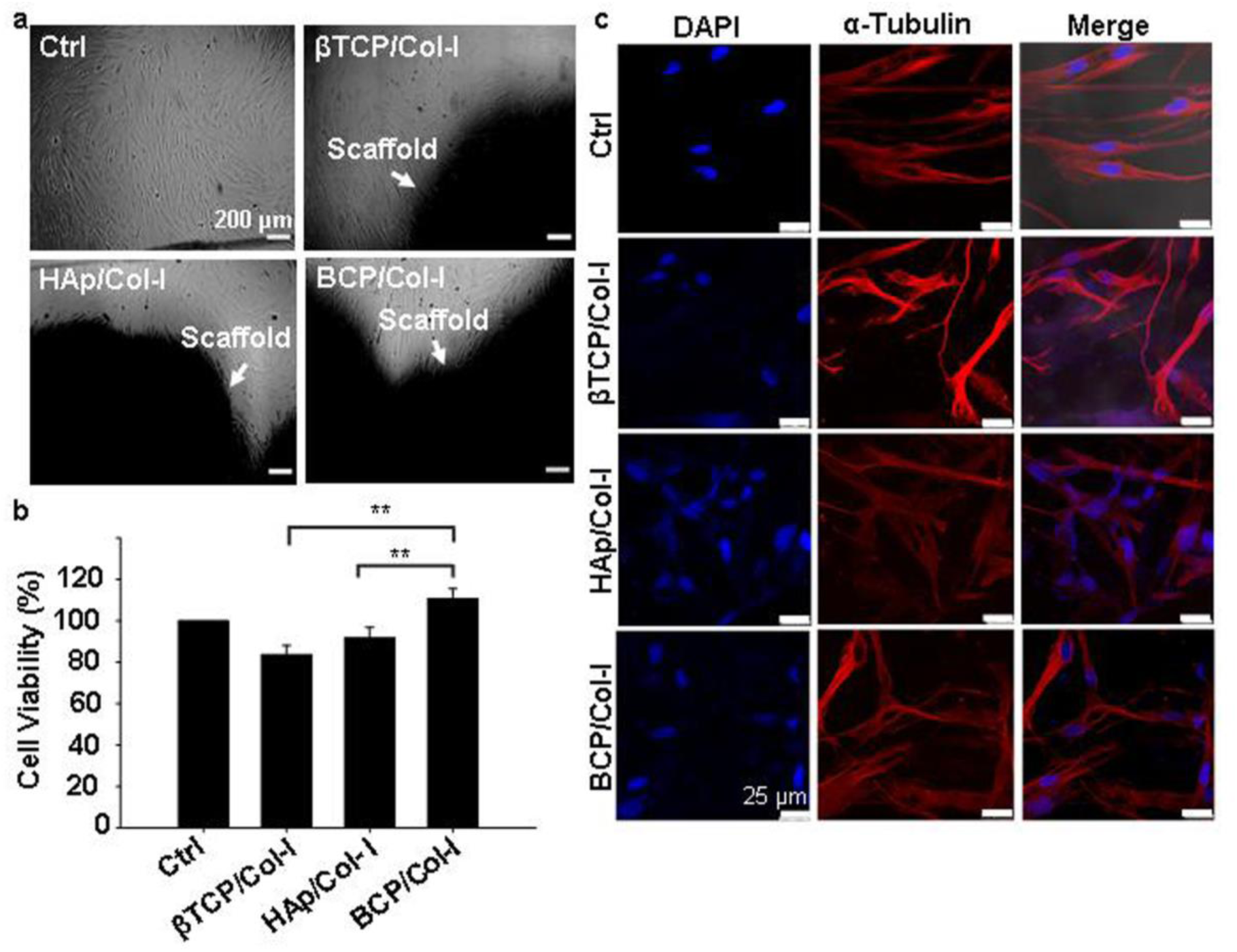
*In vitro* biocompatibility test of CP/Col-I 3D scaffolds. (a) phase contrast microscope images of g-MSCs cultured in presence of scaffold for 72 h; (b) quantitative analysis of cell viability determined by MTT assay showed > 80% cell viability for all the three CP/Col-I composite scaffolds suggesting the biocompatible nature of scaffolds with gMSCs. Histogram represents the percentage cell viability with respect to the control cells (Ctrl, 100%, n=3; mean ± SEM, \**p* ≤ 0.05); (c) Immunofluorescence images of g-MSC seeded on CP/Col-I 3D scaffolds for 72 h indicated that the 3D scaffolds provided a favourable microstructure for cell attachment. The cytoskeleton and cell nuclei were stained with α-tubulin (red) and DAPI (blue), respectively.

To investigate the influence of microstructure and chemical composition of CP/Col-I scaffold on cell adhesion, gMSCs were seeded on the CP/Col-I scaffolds and maintained in culture for 3 days. Fig. 5c shows the efficacy of gMSCs adhesion and growth following their seeding on the 3D composite scaffolds. The immuno-fluorescence images for α-tubulin (cytoskeletal marker) and DAPI (nuclear marker) revealed that the 3D scaffolds provided a favourable microstructure for cell attachment and proliferation. As expected, the growth of gMSCs was in a monolayer in 2D culture i.e., in the control group, whereas the infiltration of gMSCs into the 3D scaffolds was confirmed from multilayer arrangement. Overall, the microstructure of the CP/Col-I 3D composite scaffolds supported gMSCs adhesion and proliferation without imparting any cytotoxic effects.

#### 3.2.3. Evaluation of osteogenic differentiation potential of gMSCs seeded on CP/Col-I 3D scaffolds

We have already reported the ability of gMSCs to proliferate and differentiate towards osteocyte lineage.^27^ Hence, the same was evaluated following their seeding on the developed 3D scaffolds. As shown in Fig 6a, gMSCs cultured in the maintenance medium alone did not show the calcium deposition nodules following their staining with ARS. On the contrary, gMSC cultured using the osteoinduction medium stained strongly with ARS, suggesting the validation of osteoinduction. In fact, the gMSCs seeded onto 3D scaffolds and subjected to osteoinduction could demonstrate enhanced mineralization of the extracellular matrix compared to the control. Further, a quantitative analysis of OCN confirmed the same (Fig. 6b). Moreover, among the cell-laden 3D scaffolds, the BCP/Col-I scaffold displayed significantly higher osteogenic potential than the βTCP/Col-I and HAp/Col-I scaffolds. Hence, to assess the temporal calcium ion release behaviour, the scaffolds were incubated in PBS for different time points and the rate of calcium release was monitored over time (Fig. 6c). Interestingly, the rate of calcium ion release followed an exponential pattern, and reached a saturation point after 15^th^ day. Among the CP/Col-I composite scaffolds, the calcium ion release rate from βTCP/Col-I was significantly higher than HAp/Col-I and BCP/Col-I scaffolds. The maximum concentration of Ca^2+^ release was estimated as ∼20 mM, ∼16 mM, and ∼17 mM for βTCP/Col-I, HAp-Col-I, and BCP/Col-I composite scaffolds respectively. Further, we investigated the influence of calcium ions released from the 3D scaffold potentiating osteogenic differentiation of gMSCs. Accordingly, the scaffolds were incubated in MSCs’ maintenance medium and the conditioned media (CM) from each scaffold were collected individually at various time intervals (3, 7, and 15 days of incubation). The gMSCs were maintained with the respective CM for 3 wk and the effect of Ca^2+^ ion release in the CM in supporting osteoinduction of gMSCs was assessed by ARS staining and quantifying OCN (Fig. 6d, e). As seen in Fig. 6d, gMSCs when maintained in presence of CM could differentiate into osteogenic lineage even without exposure to the osteoinduction medium. Further quantification of OCN revealed that the CM collected on day 3 (3d-CM), irrespective of the CP/Col-I scaffold usage, supported comparatively better osteogenic differentiation of gMSCs, rather than the same collected on 7^th^ (7d-CM) and 15^th^ (15d-CM) day (Fig. 6e). Among the Cp/Col-I scaffolds, the 3d-CM collected from βTCP scaffold incubation rendered comparatively better osteoinduction potential than the other two, although not to a strikingly significant level. These results led us to hypothesize that the fabricated CP/Col-I composite scaffolds by themselves might also support osteogenesis, which was reflected by the formation of mineralized bone nodules and calcium deposits. Indeed, the ongoing *in vivo* studies (to be published elsewhere) with the stated implants have further strengthened the findings presented here.

**Fig. 6:**
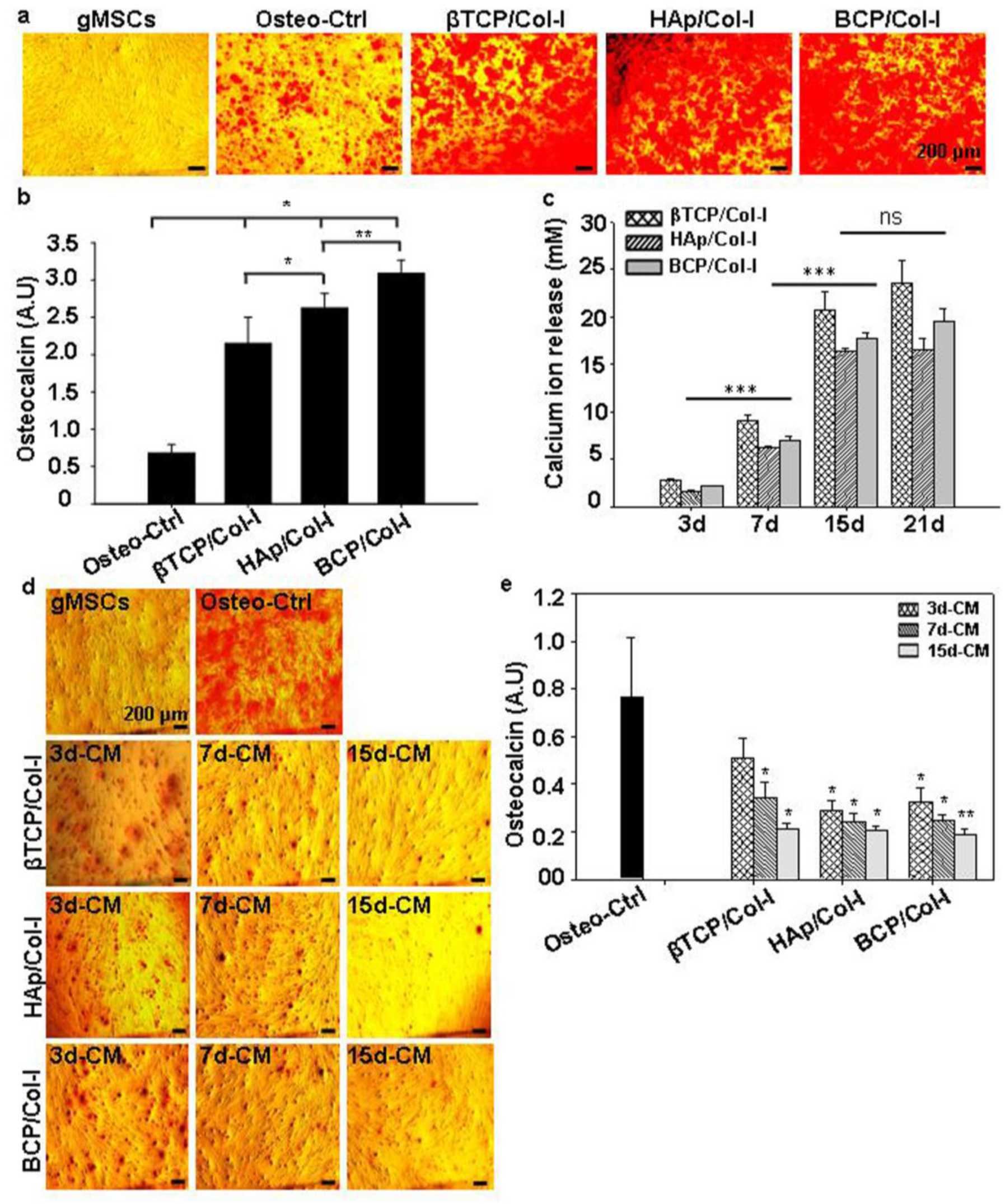
(a) Alizarin red-S (ARS) staining showed large calcium ion nodules in all 3D scaffold groups on day 21 of osteogenic induction in gMSCs. (b) Quantitative analysis of calcium ion nodules by biochemical assay demonstrated enhanced osteogenic potential of gMSCs seeded on 3D scaffolds (BCP/Col-I > HAp/Col-I > βTCP/Col-I) compared to the control (Ctrl). (c) Release of calcium ions from 3D scaffolds as a function of time (3, 7, 15, and 21 days) analysed by the EDTA titration method showed that the rate of calcium ion release from 3D scaffolds followed an exponential pattern and reached a saturation point after the 15^th^ day. (d) Microscopic images of ARS-stained MSCs that were cultured for 21 days in conditioned media (CM) prepared by incubating the CP/Col-I scaffolds for 3, 7, and 15 days in the maintenance medium. (e) Quantitative analysis of osteocalcin nodules deposited on MSCs after 21 days of culture in presence of CM of respective scaffolds for different time points revealed CM collected on day 3 (3d-CM) supported comparatively better osteogenic differentiation of gMSCs than the CM collected on 7^th^ (7d-CM) and 15^th^ (15d-CM) day. (n=4; mean ± SEM, *p ≤ 0.05).

## 4. Discussion

Bone comprises mainly of CP nanomaterial and Col-I natural polymer, and both are the most potent osteoconductive component of the extracellular bone matrix. Thus, a synthetic bone graft developed with such components is more likely to behave similarly to native bone tissue after implantation. Indeed, it is ideal to utilize CP nanomaterial and Col-I composite for the fabrication of a 3D scaffold. The 3D scaffolding methods such as stereolithography, 3D printing, electrospinning, solvent casting, and gas foaming are the most mature technologies available for fabrication. The concerns however with these methods for the fabrication of 3D scaffolds are scalability, reproducibility of microstructure, use of additives, and cost.^45^ The freeze-drying technique is currently a leading method for the fabrication of homogeneous and highly porous 3D scaffolds, although this technique involves longer processing time and requires consequential modifications for optimizing the same. However, the architecture of the 3D scaffolds by this technique includes interconnected micrometer-sized pores and porosity, which is ideal to integrate those with surrounding tissues in the body.

CP biomaterials are classified according to their Ca/P ratio and crystal structure and can have an impact on their biological properties.^5^ Typically, the HAp phase is stable, whereas the other CP nanomaterials are resorbable under physiological conditions. The CP nanomaterials resorption rate primarily depends on the Ca/P ratio; the higher the ratio, the slower becomes the resorbtion rate.^6^ Accordingly, the βTCP phase of CP nanomaterials shows biodegradability and higher resorbtion property when compared with the HAp phase, which is advantageous for new bone formation.^6^ However, the solubility or resorbtion rate of HAp phase can be altered by combining the βTCP phase into its structure, resulting in biphasic CP nanomaterial. In fact, the pH alteration shown by us in addition to the Ca/P molar ratio can also influence the synthesis of CP nanomaterials. Earlier reports have suggested that using Ca/P molar ratio at 1.5 and varying either the pH (5.5 - 8.5) or the sintering temperature (> 650 °C) can yield the βTCP phase.^46-48^ Likewise, when Ca/P molar ratio is maintained at 1.67 and sintering at 1000 °C, the HAp phase can be obtained at pH-10 and above.^33^ However, Khiri et al. have reported that the βTCP phase starts appearing by keeping the sintering temperature between 1200 – 1400 °C and that eventually forms the BCP phase due to the thermal decomposition of the HAp phase.^49^ In contrast to their findings, we have successfully obtained all three phases of CP that include BCP too, by keeping the constant Ca/P molar ratio at 1.67 and maintaining uniform sintering temperature at 1000 °C, but with pH (βTCP: 8; BCP:8.5 -9.5; HAp: 11) as the only variable. Wang et al. have also demonstrated the influence of pH (8 - 11) on different phase formations of Ca/P.^50^ While they have kept the initial precursor Ca/P molar ratio at 1.67 similar to ours, the sintering temperature for them was at 700 °C, unlike that of 1000 °C that we have used. Moreover, they have reported the pH-9 maintained samples yielding weak βTCP in HAp and with poor crystallinity, unlike distinct βTCP and HAp biphasic patterns with better crystallinity observed by us. The variation in the peak intensity and crystallinity in BCP might be due to the low sintering temperature used by them. As expected, the conjugation of Col-I with CP in the fabricated CP/Col-I scaffolds displayed broadened XRD peaks with reduced crystallinity. In fact, similar findings have been reported by Zhou et al and Asimeng et al during the fabrication of HAp/Col I composite scaffold.^51,52^

Bone is a complex tissue that provides mechanical strength to the body. During normal functioning, the bone endures bending, stretching, torsion, and compression against applied forces. The mechanical property of a CP-based composite scaffold majorly relies on the interaction between CP and the polymer. Besides, the appropriate selection of composite material is the prerequisite for the long-term reliability of 3D scaffolds as bone implants, which relies on the understanding of the biological response of the scaffold material. In fact, flexural strength and structural integrity describe the resistance of scaffolds in both tensile and compression share forces. However, the compressive strength of HAp/βTCP scaffolds varies between 3 - 50 MPa for 80 and 25% porous scaffolds.^53-55^ These values are either equal to or higher than that of human cancellous bone (2-12 MPa), where the tensile strength of cancellous bone is 2/3rd of the compression strength. Indeed, the tensile strength of CP/Col-I scaffolds fabricated by us matched well with the tensile strength of a trabecular bone (1-5 MPa).^43^

Monitoring the size of nucleated apatite globules on the surface of scaffolds is important in order to understand the effect of Ca-P composition on bioactivity. Li et al. have demonstrated that calcium-deficient HAp promotes bone-like apatite nucleation on the surface of the scaffolds.^56^ In line with the same, we have noticed a superior nucleation rate on BCP/Col-I composite scaffold compared to HAp/Col-I and with the least nucleation seen in the case of βTCP/Col-I. The presence of the bioresorbable CP phase might have contributed to the increase in nucleation rate in BCP/Col-I; as the resorbtion rate of CP nanomaterials influences the apatite nucleation on the surface of CP scaffolds.^57^ Moreover, the nucleation of the apatite layer could be triggered by providing binding sites for calcium and phosphate ions from cationic species such as -NH_2_^2+^ from Col-I and anionic species such as PO_4_^3-^ from 3D composite scaffolds.^58^

In the current state-of-the-art scaffold-based bone tissue engineering research, attempts have been made to develop artificial 3D scaffolds with a combination of cells that would mimic both the composition and structure of bone. The chemical composition, surface architecture, and pore geometry, including porosity and pore interconnectivity of the scaffold are crucial in regulating cellular behaviour.^59^ Hence, it is generally agreed that a highly porous 3D scaffold coupled with interconnected pores and a large surface area is favourable for tissue in-growth.^60^ Moreover, the cell viability and adhesion to the surface of the synthetic bone scaffold are the key factors in bone tissue engineering. Similarly, the pore size of the scaffolds may also carry a significant attribute in cell adhesion. While Murphy and O’Brien have investigated the microporous 3D scaffolds for bone tissue engineering using MC3T3-E1 cells, Zhang et al. have reported the direct correlation of optimum pore size (100-300 μm) with MSCs growth.^59,61^ In line with the same, the CP/Col-I 3D composite scaffolds fabricated by us by freeze-drying had carried the pore dimension ranging from 80-450µm and had supported efficient gMSCs adhesion and growth. Wei et al. have stated that the porous 3D scaffold with enhanced surface area promotes cell attachment and proliferation.^62^ In fact; our data has suggested that the microstructure of the CP/Col-I 3D composite scaffold may be favourable for gMSCs adhesion and proliferation without imparting any cytotoxic effects. The higher surface area and porosity of scaffolds might have boosted protein adsorption, bioactivity, and other biological activities, *albeit* with a predominant trade-off between the porosity and mechanical strength of the scaffolds.^26,43^ Among the scaffolds BCP/Col-I having higher surface area, reasonable porosity, and mechanical strength was found to be the most suitable one for gMSCs growth and osteogenic differentiation.

A number of reports indeed suggest that calcium ion released from bone resorbtion surfaces promotes cell proliferation and matrix mineralization of MSCs.^63,64^ Moreover, another clue comes from the fact that CP nanomaterials tend to release Ca^2+^ ions from their structure.^65^ However, the influence of calcium ion leached from CP nanomaterial on MSCs’ differentiation toward osteocyte lineage both *in vitro* and *in vivo* remains unclear. Maeno et al. have shown that 2-4 mM Ca^2+^ ion is suitable for proliferation and survival, whereas 6-8 mM Ca^2+^ ion can favour osteoblast differentiation and matrix mineralization of primary osteoblasts cells.^66^ Further, Jung et al. have demonstrated that the Ca^2+^ ion released from the HAp structure can accelerate the osteoblast differentiation of MC3T3-E1 cells.^67^ However, our findings suggested that even as low as 3 mM Ca^2+^ ion release from the scaffold can induce osteogenic differentiation from gMSCs.

## 5. Conclusion

Synthetic bone grafts with extracellular matrix (ECM) composition provide the ideal platform for cell adhesion and stem cell differentiation to bone tissues. Although ceramic materials have been the ideal choice for the development of scaffolds to enhance the effectiveness of bone grafts, of late, composite materials with ceramic-polymer components have drawn immense attention for bone tissue engineering applications. Herein, we have systematically investigated the osteo-differentiation potential of gMSCs seeded on different phases of CP nanomaterials in conjunction with Col-I 3D assembly. We were primarily interested in investigating the impact of different Ca/P ratios on microstructural, physicochemical, and mechanical properties of freeze-dried CP/Col-I 3D scaffolds, and their associated impact on *in vitro* biological properties for bone graft application. Indeed, gMSCs having the multipotent differentiation potential and intrinsic ability to differentiate into osteocytes could serve as the ideal cell of choice to use these in combination with the afore-stated nanocomposite scaffolds and explore their efficacy in promoting osteoinduction. Among the scaffolds, the BCP-Col-I with better microarchitecture, high surface area combined with an adequate tensile strength similar to that of trabecular bone has provided the impetus for their further investigation *in vivo* using animal models with bone defects. Moreover, the association of calcium ion release from the fabricated scaffolds with osteoinduction efficacy holds promise for their further exploration in bone tissue regeneration as a cell-free approach.

## 6. Conflict of Interest

There is no conflict of interest to declare.

## 7. Acknowledgements

NL thanks the extramural funding support from NER-BPMC, Dept. of Biotechnology, Government of India (BT/PR16655/NER/95/132/2015) and intramural funding received from NCCS. MMB is a WOS-A fellow supported by the Department of Science and Technology (DST), Govt. of India (SR/WOS-A/LS-602/2016). AR is an N-PDF supported by DST (PDF/2020/002202). The funders have no role to play in the study design, data collection, analysis and interpretation of the findings and submission of the manuscript for publication.

## 8. Author Contributions

MB, AR, and NL actively participated in the collection and/or assembly, analysis, and interpretation of data, and in writing the manuscript; DP and NL supervised the research and all were involved in the final approval of the manuscript.

## 9. Supporting Information

**Fig. S1.** Original FTIR spectra of CP nanomaterials synthesized at pH 8, 9, and 11, respectively.

**Fig. S2.** (a, b): EDAX and FE-SEM of CP/Col-I scaffolds after 32 days of incubation; a) atomic % and weight % elemental present in the newly formed apatite, b) EDAX spectra and FE-SEM micrographs.

## Notes

### Competing Interest Statement

The authors have declared no competing interest.

