## Supplemental Figures S1 and S2 for "Fabrication and Characterization of Ceramic-Polymer composite 3D scaffolds and Demonstration of Osteoinductive propensity with gingival Mesenchymal Stem Cells"

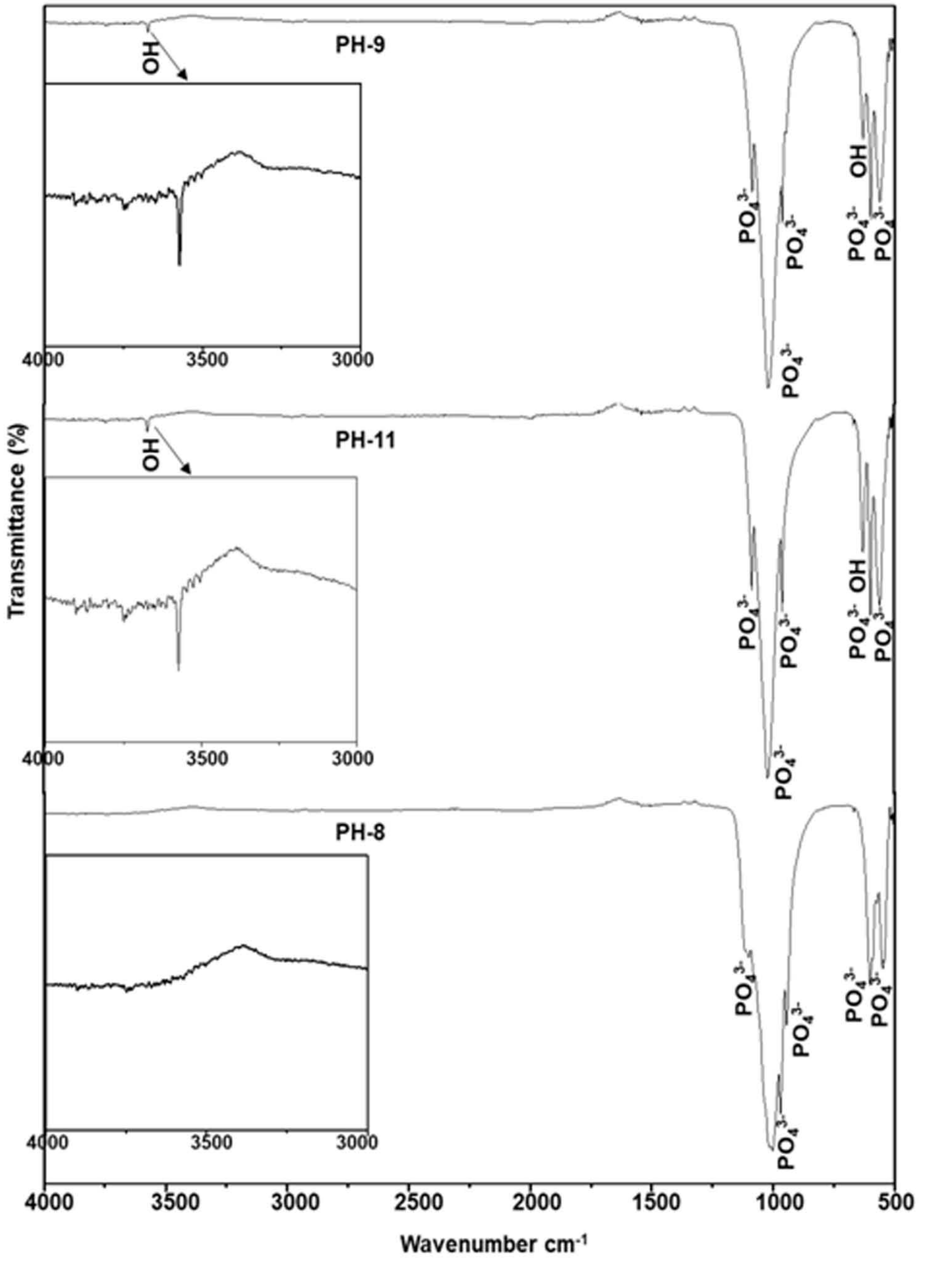


**Fig. S1.** Original FTIR spectra of CP nanomaterials synthesized at pH 8, 9 and 11 respectively.

**a**

| Element | Weight% | Atomic% |
|---|---|---|
| O K | 43.20 | 63.20 |
| Na K | 2.49 | 2.53 |
| Mg K | 0.25 | 0.24 |
| P K | 14.27 | 10.79 |
| K K | 0.20 | 0.12 |
| Ca K | 39.59 | 23.12 |
| Totals | 100.00 | |

| Element | Weight% | Atomic% |
|---|---|---|
| O K | 51.80 | 69.46 |
| Na K | 4.92 | 4.59 |
| Mg K | 0.80 | 0.70 |
| P K | 15.84 | 10.97 |
| K K | 0.26 | 0.14 |
| Ca K | 26.38 | 14.12 |
| Totals | 100.00 | |

| Element | Weight% | Atomic% |
|---|---|---|
| O K | 40.31 | 59.74 |
| Na K | 2.36 | 2.44 |
| Mg K | 0.67 | 0.65 |
| P K | 20.94 | 16.03 |
| K K | 0.27 | 0.16 |
| Ca K | 35.45 | 20.97 |
| Totals | 100.00 | |

**b**

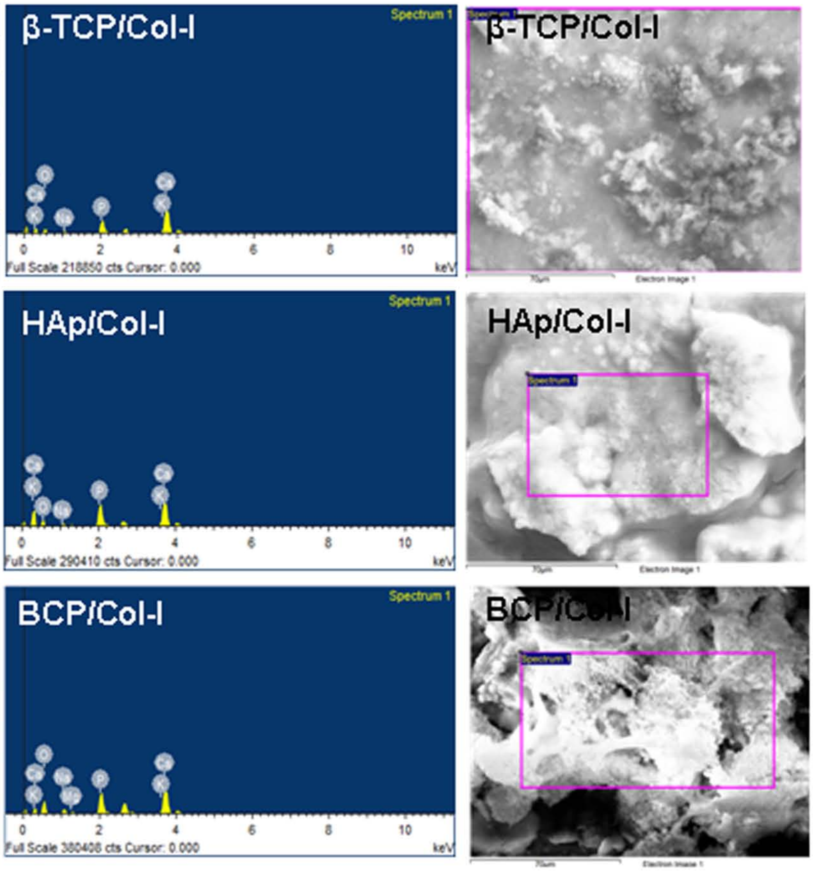


**Fig. S2** (a, b). EDAX and FE-SEM of CP/Col-I scaffolds after 32 days of incubation; a) atomic % and weight % elemental present in the newly formed apatite, b) EDAX spectra and FE-SEM micrographs.
